# Genotoxic stress and clonal expansion shape the somatic genome of systemic sclerosis skin

**DOI:** 10.64898/2026.11.26.769567

**Authors:** Robert Gniadecki, Dylan Hennessey, Sandra O’Keefe, Junqin Wang, Jan Storek, Caylib Durand, Aishwarya Iyer, Mohammed Osman

## Abstract

1.

**Background:** Systemic sclerosis (SSc) is a life-limiting inflammatory disease characterised by skin and visceral fibrosis. We previously showed that fibrotic skin from patients with diffuse cutaneous SSc (dcSSc) accumulates a large number of somatic mutations, although the mechanisms that generate them and their disease-related consequences remain poorly understood. We investigated the burden, clonal architecture and biological significance of somatic mutations in dermal sections from patients with dcSSc (<2 years from the first non-Raynaud’s symptom), dcSSc treated with autologous haematopoietic stem cell transplantation (post-AHSCT) and limited cutaneous SSc (lcSSc, <5 years from the first non-Raynaud’s symptom), together with dermal fibroblasts (DFs) from each group.

**Methods:** We performed whole exome sequencing of 52 lesional skin biopsies and 33 low-passage (passage 2) DF lines with autologous buccal DNA as the germline reference. Dermal samples comprised 20 dcSSc, 22 lcSSc and 10 post-AHSCT biopsies, and 21 DF from dcSSc patients, and 12 in the post-AHSCT group. Mutational burden, signatures and variant allele fractions were related to clinical phenotype, fibroblast transcriptomes and replication timing, and post-AHSCT samples were used to assess persistence of mutant clones.

**Results:** Mutations were carried by clonally expanded dermal cell populations, and represented a median of 17% of sampled cells, without evidence of positive selection. A large component of all mutations (>25 %) had a sequence linked with oxidative guanine damage (C>A), pointing to ongoing DNA damage and cellular stress. The C>A burden was higher in dcSSc than in lcSSc and correlated with the modified Rodnan skin score - a clinical measurement of skin fibrosis severity. Variant burden increased with patient age but not with disease duration. AHSCT did not change the mutational landscape.

**Conclusions:** Fibrosis skin in SSc is characterized by a widespread somatic mosaicism generated by ongoing mutagenesis attributable to oxidative DNA damage. Persistence of mutant clones after regression of fibrosis argues against their being sufficient drivers of the fibrotic phenotype and identifies the somatic genome as a record of cellular injury.

## 2. Introduction

Systemic sclerosis (SSc) is generally viewed as an immune-mediated fibrotic disease in which vascular injury, inflammation and persistent fibroblast activation drive progressive tissue fibrosis, resulting in life-limiting complications and severe disability [1]. This mechanism, however, has largely overlooked acquired genetic alterations within the affected tissues. We previously sequenced microdissected fibrotic dermis from patients with rapidly progressive diffuse cutaneous SSc (dcSSc) using autologous buccal DNA as the germline reference and found an increased burden of somatic mutations with a predominantly clock-like substitution pattern [2]. An increased somatic mutation burden was subsequently reported in cultured fibroblasts from SSc lung, with signatures attributed to translesion polymerase and deaminase activity [3]. Somatic mutagenesis has therefore been demonstrated in fibrotic SSc tissue from two organs using independent approaches, but its origins and biological significance remain uncertain.

Oxidative stress is one plausible source. Dermal fibroblasts (DFs) from affected skin constitutively generate reactive oxygen species through NADPH oxidase, which helps maintain their activated phenotype in culture [4], and urinary 8-hydroxy-2′-deoxyguanosine, an indicator of oxidative DNA damage, is increased in SSc and correlates with the modified Rodnan skin score in dcSSc [5]. Mitochondrial dysfunction may provide an additional source. We recently found that DFs from patients with dcSSc carry increased mitochondrial and total cellular reactive oxygen species, increased markers of double-strand DNA breaks, senescence-associated signals and resistance to apoptosis [6], and that fibrotic dermal niches in SSc show indicators of mitochondrial dysfunction [7]. DNA damage signals and chromosomal instability in lesional fibroblasts have been linked to cGAS-STING activation [8] and to senescence-associated signals [9–11].

Somatic mutations may contribute directly to disease by providing a proliferative advantage to fibroblasts, analogous to the expansion of driver-mutant clones in normal epithelia [12,13]. The recurrent mutation of cancer-associated genes, including *KMT2C*, *DMD* and *ARID1A*, found in SSc skin is compatible with this possibility [2], but recurrent mutation alone does not establish selection because mutation frequency is strongly influenced by gene size and local mutability. Other observations argue against a simple cancer-like driver model. Somatic mutation in SSc fibroblasts is not accompanied by mesenchymal transformation or overt hyperplasia, and the increased cancer risk in SSc is concentrated in lung, liver, haematological and bladder malignancies rather than connective-tissue tumours [14].

In this study we sequenced lesional skin and cultured DF exomes from patients with dcSSc (<2 years from the first non-Raynaud’s symptom), patients with dcSSc treated with autologous haematopoietic stem cell transplantation (post-AHSCT) and from patients with limited cutaneous SSc (lcSSc, <5 years from the first non-Raynaud’s symptom), using autologous germline DNA as the reference. We integrated variant allele fractions with clinical data, opportunity-corrected models of selection and fibroblast transcriptomes, and found that somatic variants were abundant and carried by expanded clones that showed no evidence of positive selection. The oxidative mutational signature was associated with SSc subtype and with accumulated skin fibrosis burden, and mutant alleles remained detectable after AHSCT. Together, these observations suggest that mutant clones are by themselves insufficient to drive fibrosis and that the mutations are better read as indicators of ongoing cellular injury.

## 3. Results

### 3.1. SSc skin carries a C>A substitution component of oxidative type

To characterise somatic variation in SSc, we sequenced exomes from 52 lesional skin samples from 48 patients and 33 cultured DF samples from 29 patients using autologous buccal DNA as the germline reference; 22 patients contributed both tissues. Skin was microdissected from fibrotic dermis as we have described before [2] to minimise the ultraviolet-associated signal that dominates whole-skin biopsies and originates largely from mutated epidermal clones [12]. Calls supported by only one or two reads were further assessed for sequencing error. Hence, our primary analyses required at least three alternate reads and were repeated across increasing levels of read support. At this threshold, lesional skin and cultured SSc fibroblasts contained a median of 245 private coding variants per exome, corresponding to 22.7 variants per callable megabase (Fig. 1a,b).

**Figure 1.**
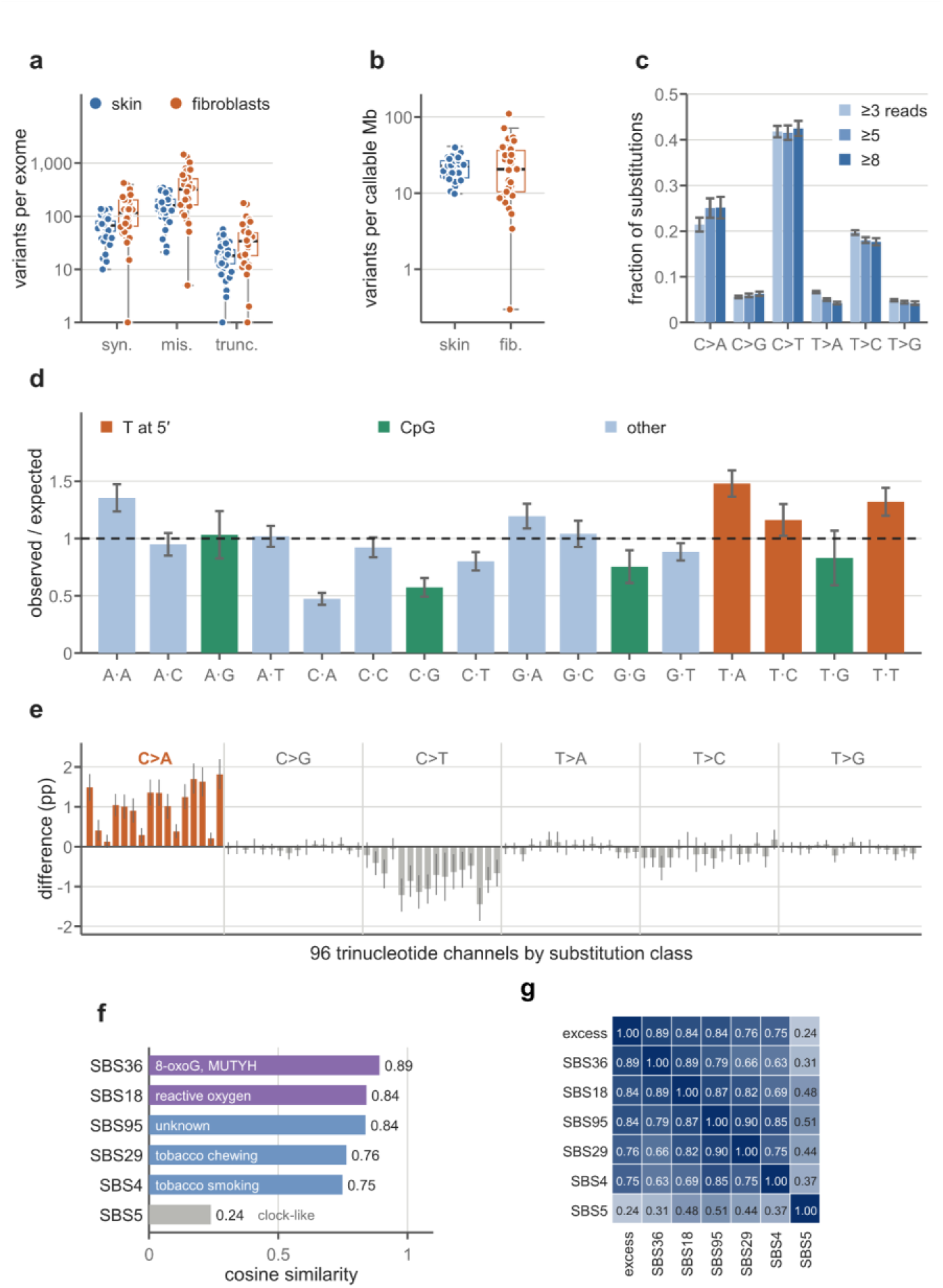
Somatic variants in SSc skin carry a C>A component with an oxidative-type sequence context. Private coding variants from 52 lesional skin exomes (48 patients) and 33 cultured dermal fibroblast exomes (29 patients), called against autologous buccal DNA and supported by at least three alternate reads unless stated. (a) Variants per exome by predicted consequence. (b) Variants per callable megabase. (c) Six-class spectrum of skin substitutions at three, five and eight or more alternate reads, averaged within and then across 42 patients (mean ± SE) (d) Trinucleotide context of skin C>A substitutions at eight or more reads (27 patients) as observed over expected share, the expectation taken from the trinucleotide composition of the canonical coding sequence (mean ± SE across patients). (e) Difference between the 96-channel spectra of the 23 skin exomes with the highest and the 23 with the lowest C>A fraction. (f) Cosine similarity of the positive part of the excess spectrum to exome-scaled COSMIC v3.4 signatures. (g) Pairwise cosine similarity among the excess spectrum and the same signatures. Boxes span the interquartile range with the median, whiskers 1.5 times the interquartile range; points are exomes.

C>T transitions were the most frequent substitutions, while C>A transversions consistently accounted for approximately one fifth to one quarter of substitutions in skin (Fig. 1c). C>A substitutions can arise when 8-oxoguanine (8-oxoG) is generated by oxidative DNA damage and then mispairs with adenine during replication, but they can also result from bulky DNA adducts such as those produced by tobacco carcinogens [15]. The pattern in SSc skin favoured the former explanation: C>A substitutions occurred equally on the transcribed and non-transcribed strands, were depleted rather than enriched at CpG dinucleotides and were enriched when preceded by thymine (Fig. 1d), and they occurred at similar frequencies in never-smokers and in patients with a history of smoking. Together, these features are inconsistent with a helix-distorting adduct and more compatible with mispairing of 8-oxoG.

We next asked whether the C>A component corresponded to a recognised mutational signature. Fitting reference signatures to the complete spectrum was uninformative because the clock-like signature SBS5 accounted for most substitutions, while the C>A component was assigned to SBS4, SBS29 or SBS36 depending on the reference set. We therefore isolated the component that differed most between patients by subtracting the mean spectrum of the exomes with the lowest C>A fraction from that of the exomes with the highest. The resulting excess consisted almost entirely of C>A substitutions in thymine-preceded contexts (Fig. 1e). This spectrum resembled several C>A-rich reference signatures of high degree of similarity which precluded its assignment to a single reference mutational process (Fig. 1f, g). In summary, somatic variation in SSc skin contains a reproducible C>A component whose sequence context points to oxidative guanine damage without identifying a single catalogued process.

### 3.2. The C>A fraction tracks subtype and accumulated skin disease

dcSSc is characterised by more extensive skin involvement, higher modified Rodnan skin scores (mRSS) and greater internal-organ involvement than limited cutaneous disease [16,17], and progression of skin fibrosis predicts declining lung function and worse survival [18]. We therefore asked whether the somatic mutational landscape differed between the two subtypes. Overall variant burden was not associated with subtype or skin score. In contrast, the proportion of C>A substitutions was markedly higher in dcSSc than in lcSSc, and the difference became more pronounced as weakly supported calls were excluded and remained after adjustment for sequencing depth and call number (Fig. 2a).

**Figure 2.**
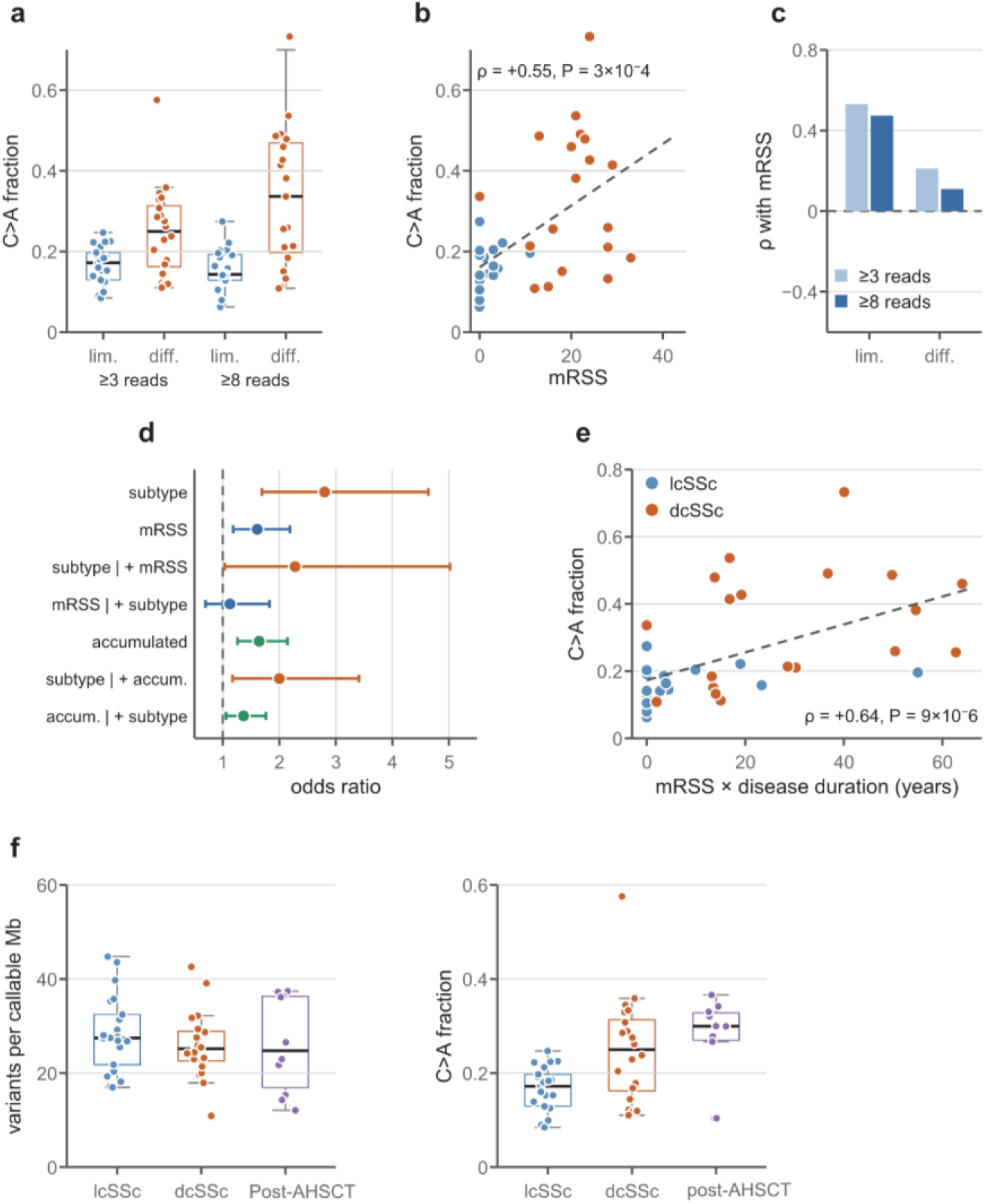
The C>A fraction tracks subtype and accumulated skin disease. (a) C>A fraction of single-nucleotide substitutions by subtype at three (41 exomes) and eight or more alternate reads (39 exomes with an mRSS). *Significant odds ratios of 1.90 (95% CI 1.32–2.74) and 2.80 (CI 1.69–4.64). (b) C>A fraction at eight or more reads against mRSS, with least-squares fit. (c) Spearman correlation of mRSS with C>A fraction within each subtype; early limited ρ = 0.53 (P = 0.016) and 0.47 (P = 0.034), early diffuse 0.21 and 0.11 (both P > 0.3), at three and eight reads. (d) Odds ratios (95% CI) from binomial models of C>A against non-C>A counts with patient-clustered errors, at eight or more reads, according to disease subtype and accumulated damage (mRSS × disease duration). (e) C>A fraction at eight or more reads against mRSS × disease duration (ρ = 0.64, P = 9 × 10⁻⁶); exomes with a zero score or duration have an index of zero. (f) Effect of AHSCT, data for 10 dermal and 12 fibroblast exomes from 11 patients, 2 to 33 months after transplantation. Coding variant density (left) and C>A fraction (right) of untreated lcSSc, dcSSc, and post-transplant dcSSc skin.

The C>A fraction also increased with mRSS (Fig. 2b), an association that was more apparent within lcSSc but not within dcSSc (Fig. 2c). Because subtype and mRSS are strongly correlated, we modelled them together. At the more stringent read-support threshold, subtype remained associated with the C>A fraction whereas current mRSS did not, and at the primary threshold neither remained independently significant (Fig. 2d). Because mRSS captures current skin involvement whereas the mutational spectrum may integrate a longer history of tissue injury, we examined mRSS multiplied by disease duration as an index of accumulated skin disease. This index correlated with the C>A fraction (Fig. 2e) and remained associated with it after adjustment for subtype, with both terms stable in leave-one-patient-out analyses (Fig. 2d). The oxidative C>A component therefore appeared to reflect cumulative skin disease in addition to subtype. Men had a higher C>A fraction than women, but only 13 of the 41 exomes came from male donors, which prevented reliable separation of sex from disease severity.

Finally, in a prespecified grid relating four measures of the somatic landscape to 13 clinical and serological variables, no association remained significant after correction for multiple testing. Taken together, these analyses indicate that C>A mutations are associated with diffuse disease and with accumulated skin involvement, providing further evidence for the importance of oxidative-type genotoxic stress.

### 3.3. Long-gene transcription falls and variants track late replication

To determine whether the mutational process was accompanied by ongoing DNA damage, we used the observation that endogenous DNA lesions can stall elongating RNA polymerase II, reducing transcriptional output in proportion to the amount of DNA traversed [19]. Because introns account for most of that distance, we compared fibroblasts from SSc and normal skin in the single-cell RNA sequencing dataset GSE249279 [20], as we have done previously [21], stratifying genes by intronic length while holding mature transcript length constant. In all three major fibroblast subtypes, expression in SSc declined progressively as intronic length increased (Fig. 3a), a pattern consistent with damage-associated transcriptional stalling.

**Figure 3.**
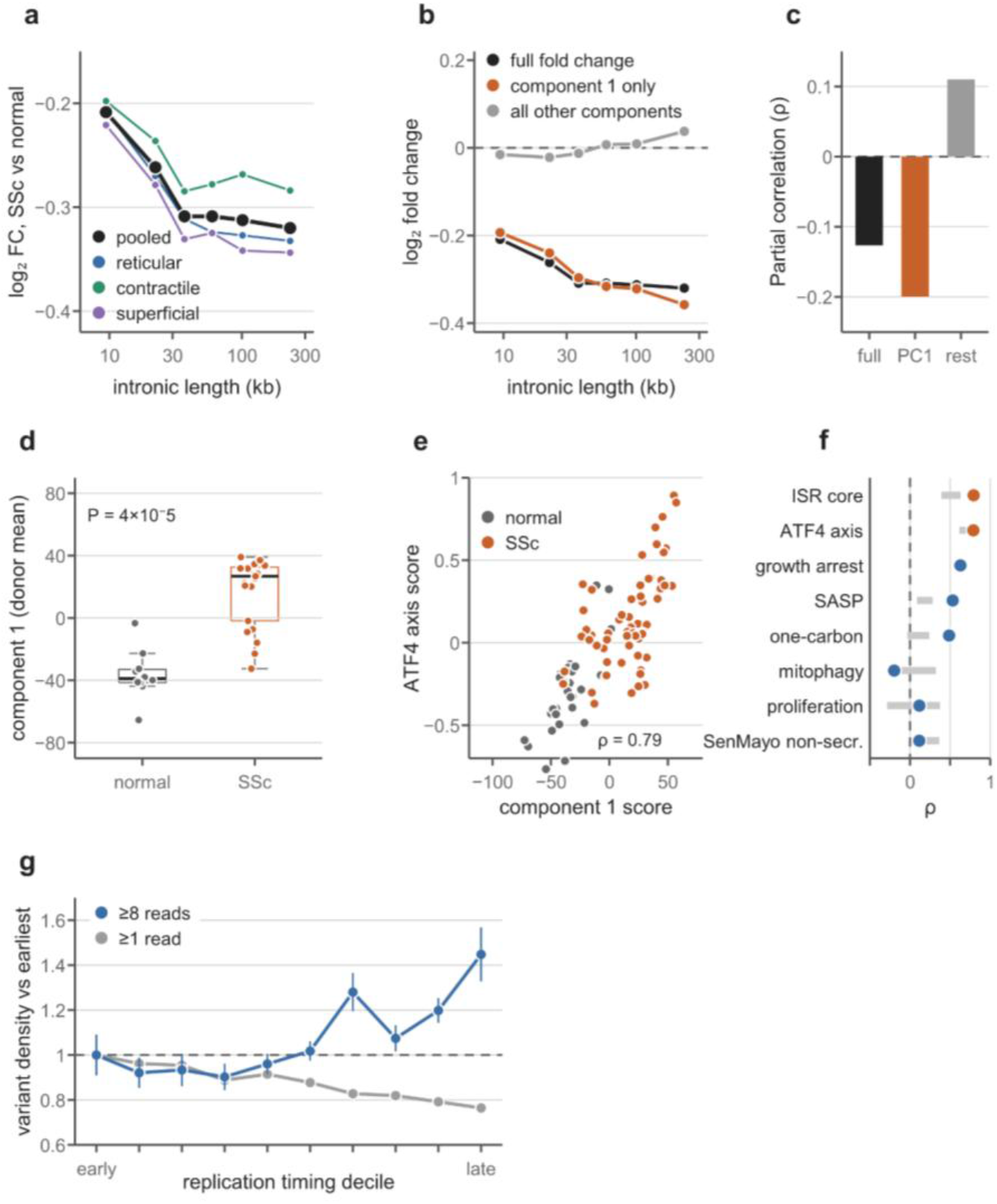
Length-dependent loss of transcription in SSc fibroblasts carried by an integrated stress response programme. Data from fibroblasts of the dataset GSE249279 [20], aggregated to pseudobulk log-CPM per donor and fibroblast subtype (reticular, contractile, superficial). (a) Mean log2 fold change in six bins of intronic length within quintiles of mature transcript length. (b) Fold change reconstructed from principal component 1 (orange) and from all other components (grey), binned as in (a). (c) Rank partial correlation of fold change with intronic length at fixed transcript length. (d) Component 1 score per donor; two-sided Mann– Whitney P = 3.8 × 10⁻⁵. (e) Component 1 against the length-matched ATF4-axis score per unit (Spearman ρ = 0.79). (f) Spearman correlation of component 1 with length-matched module scores across all units (points) and the range of the within-SSc and within-normal correlations (grey bars); ISR core 0.79 and *ATF4* axis 0.79 (both P < 10⁻⁴ after adjustment for log library size and log cell number). (g) Variant density of 27 fibroblast exomes (24 patients) by replication-timing decile of BJ fibroblasts, relative to the earliest decile, at eight or more (blue; mean ± SE across 20 patients) and one or more alternate reads (grey). The per-patient timing coefficient was negative in 19 of 20 patients.

We then asked whether this length-dependent transcriptional loss represented a diffuse property of all genes, as described in ageing tissues [22]. Principal-component decomposition of the matrix of fibroblast expression values, aggregated per donor and fibroblast subtype, showed that the first component alone reproduced the decline in expression with increasing intronic length (Fig. 3b, c), and the same component separated SSc from normal donors (Fig. 3d). The strongest correlates of this transcriptional programme were genes downstream of the ATF4-dependent integrated stress response, and this association persisted after comparison with length- and expression-matched control genes, within SSc and normal fibroblasts separately, and after adjustment for library size and cell number (Fig. 3e, f).

A similar dependence was seen for mature transcript length. Reduced transcription of long genes in SSc fibroblasts was therefore linked to the integrated stress response, consistent with persistent cellular stress [23] and with the mechanism we have recently described in these cells [6,21].

Because the human mutation rate depends on the timing of DNA replication [24], we next used the known replication-timing profile of BJ cells, a normal human foreskin fibroblast strain, as a reference [25] and compared it with the variant distribution in low-passage SSc fibroblasts cultured for fewer than two passages. Variant density was substantially higher in late-replicating than in early-replicating genes, with the same direction of effect in 19 of 20 patients (Fig. 3g), consistent with the known enrichment of substitutions in late-replicating DNA [24]. This effect was not caused by defective transcription-coupled repair, because mutation density was highest in silent genes and decreased with increasing expression, which is expected when transcription-coupled repair is preserved, and no substitution class showed strand asymmetry in expressed or silent genes. Adjustment for replication timing also removed the apparent excess in silent genes. Chromatin accessibility was not measured in these samples, so replication timing serves here as a partial proxy for the chromatin context in which variants accumulate. Thus, the mutational process in SSc fibroblasts was accompanied by a transcriptional signature of lesion-associated stalling and by the replication-linked positional bias expected of errors fixed during DNA synthesis.

### 3.4. Somatic variants in SSc skin reside in expanded clones

Because replication-associated mutagenesis (Fig. 3g) requires cell division, we asked whether mutated fibroblasts expand and give rise to cellular clones in the skin. Among private coding variants sequenced to a depth of at least 30-fold, the median variant allele fraction (VAF) was 0.092, corresponding under heterozygosity to a clone occupying approximately one fifth of the sampled cells, and fewer than 2% of variants had a VAF below 0.02 (Fig. 4a). The observed distribution therefore indicated that the detected variants arose in cells that subsequently expanded clonally.

**Figure 4.**
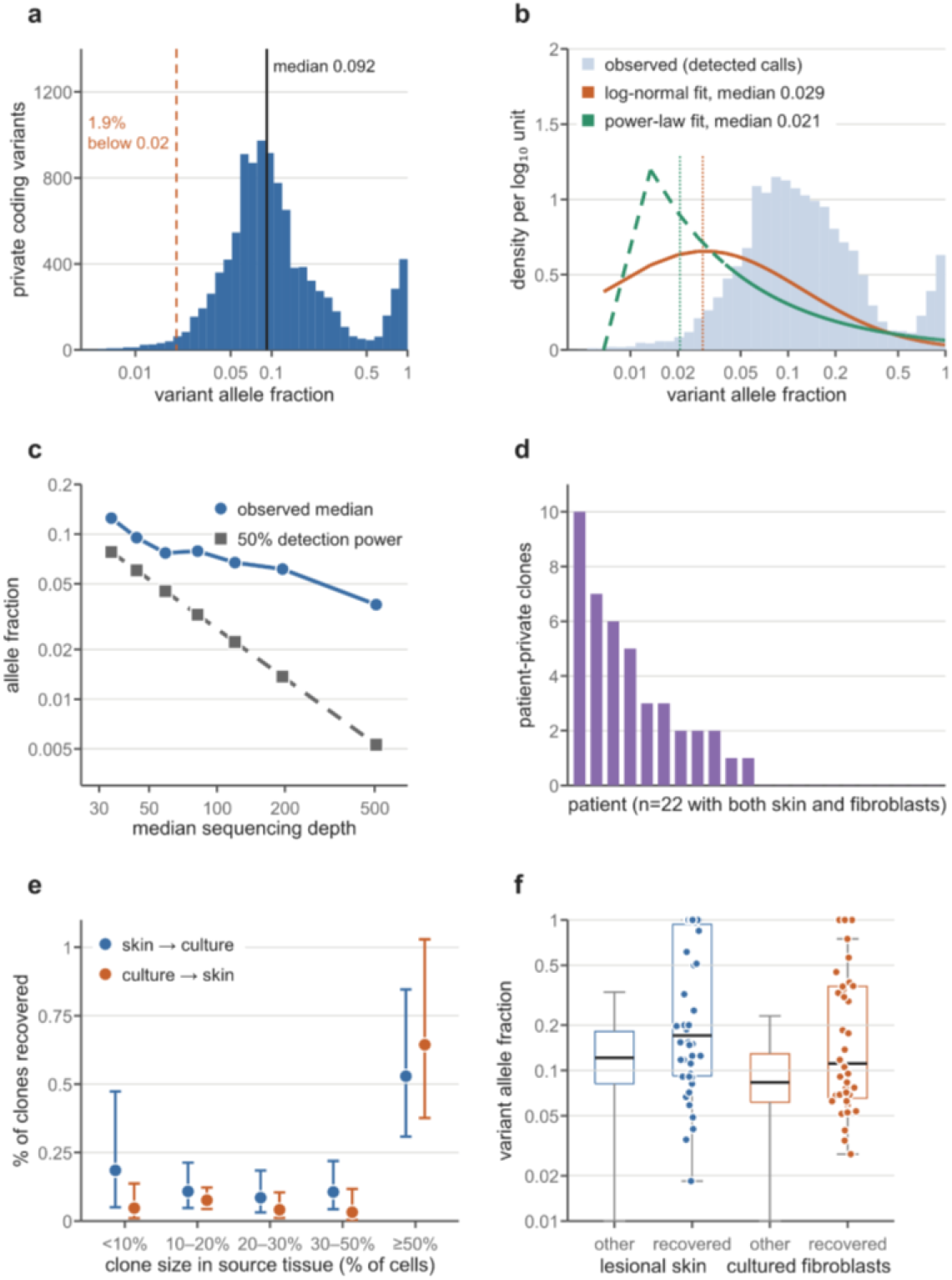
Somatic variants in SSc dermis are carried by expanded clones. Clone size is the fraction of sampled cells carrying a variant, twice the allele fraction under heterozygosity. (a) Allele fractions of private coding variants in 52 skin exomes with at least three alternate reads and 30-fold depth (n = 10,337); median 0.092 (interquartile range 0.062–0.160), 1.9% below 0.02 (dashed). (b) Allele fractions of the 15,988 calls entering the deconvolution (bars) with latent distributions estimated by maximum likelihood conditional on detection: log-normal median 0.029 (best by Akaike information criterion) and truncated power law 0.021; beta and gamma families were not identified on this call set. (c) Median observed allele fraction in seven depth strata (blue) and the allele fraction at 50% detection power (grey); 0.037 at 300-fold or more (999 calls). (d) Patient-private clones detected in both lesional skin and the fibroblast culture from the same biopsy, per patient (22 patients; 42 clones in 11). (e) Percentage of clones in one tissue recovered in the paired tissue, by clone size in the source tissue, with 95% binomial intervals; calls of any read support, of which 22 of the 42 recovered clones carry three or more reads in both tissues. Odds of recovery per 0.1 allele fraction 1.45 (95% CI 1.23–1.72) from skin to culture and 1.64 (1.33–2.01) from culture to skin (logistic models, patient-clustered errors). (f) Allele fraction of recovered and other clones; medians 0.171 against 0.121 in skin (P = 0.0013) and 0.111 against 0.083 in culture (P = 0.004; two-sided Mann–Whitney).

Because detection of a variant required at least three supporting reads, the observed VAF distribution preferentially captured larger clones. We therefore estimated the underlying clone-size distribution by maximum likelihood, modelling every call as a binomial draw of alternate reads at its own sequencing depth and conditioning each likelihood on the probability that the call would have been detected at that depth, so that clones too small to reach the three-read threshold are accounted for rather than ignored. Across alternative distributional families, the typical detectable clone was small, involving only 1 to 3% of sampled cells (Fig. 4b). Consistent with this estimate, the observed median VAF fell as sequencing depth increased and smaller clones became detectable, reaching 0.037 at sites sequenced above 300-fold (Fig. 4c). More than half of the fitted clone distribution lay below the allele fraction at which detection reached 50% power, indicating that most clones present in the tissue were probably smaller than those directly observed.

We next asked whether dermal clones could be traced into fibroblast cultures established from the same biopsy, which was possible for the 22 patients who contributed paired skin and fibroblast exomes. Of 95 variants found in both tissues from the same patient, 53 were excluded because they were concentrated in difficult-to-sequence regions. The remaining 42 patient-private variants were distributed across 11 patients (Fig. 4d), and none was shared between skin and fibroblast exomes from different patients, supporting a shared clonal origin rather than recurrent sequencing artefact.

Recovery of the same clone in both skin and culture was uncommon but strongly dependent on clone size. Clones recovered in culture were larger in the original skin sample than clones detected in skin alone (Fig. 4e, f), and the same relationship was observed when the analysis was reversed from culture to skin. Because a fibroblast culture is founded by a fraction of the cells present in the biopsy, larger tissue clones are more likely to be represented among the founders. Consistent with this, two cultures established independently from one biopsy shared predominantly their largest clones. In summary, somatic variants in SSc skin reside in expanded, mitotically active clones, of which only the largest are reliably propagated in culture.

### 3.5. Near-neutral evolution of mutated clones in SSc

To determine whether clonal expansion could be driven by particular mutations, we first examined the most frequently mutated genes in SSc skin and fibroblasts. In both tissues, the most frequently mutated genes were among the longest in the genome (Fig. 5a), as previously observed in cancer exomes [26]. Variant counts increased with callable coding length (Fig. 5b), and genes rich in coding variants were also rich in intronic variants (Fig. 5c), indicating that much of the mutational recurrence reflected the amount of exposed DNA and local mutability rather than protein-coding consequence. Because longer genes were also more completely callable, we calibrated the exposure model using 2,000 random gene sets matched for gene architecture. An uncorrected model produced substantial false-positive enrichment, whereas two models that explicitly accounted for mutational opportunity or callability remained close to the expected false-positive rate. Both corrected models were therefore used for subsequent gene-set analyses.

**Figure 5.**
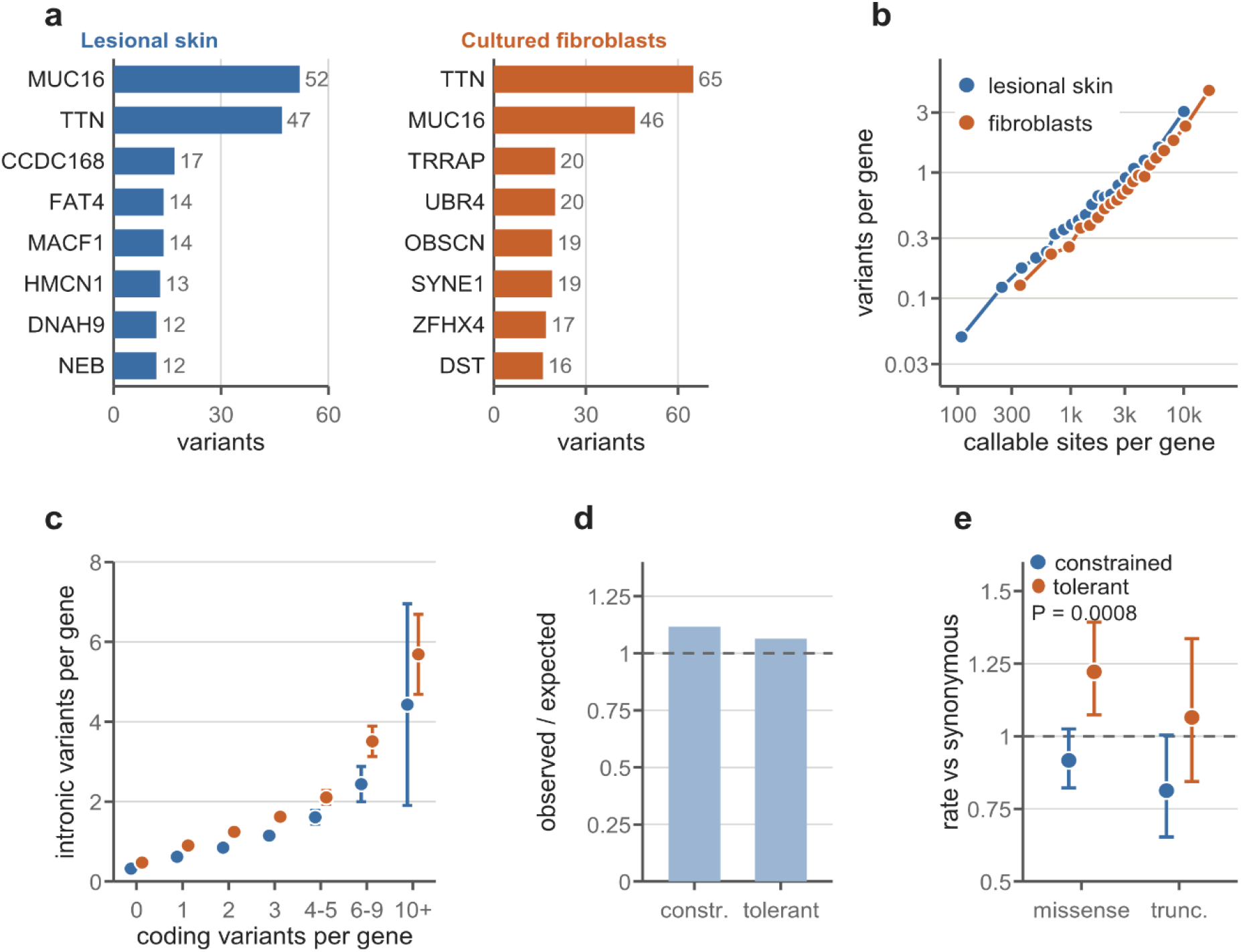
Gene mutation pattern in SSc. Private coding single-nucleotide variants with at least three alternate reads in 52 skin and 33 fibroblast exomes. (a) The eight most frequently mutated genes in each tissue, all among the longest protein-coding genes. (b) Mean variants per gene in 20 bins of callable coding opportunity, binned separately in each tissue; Spearman ρ across genes 0.46 in skin (17,791 genes) and 0.51 in fibroblasts. (c) Mean intronic variants per gene (95% CI) by number of coding variants in the same gene; ρ = 0.28 in skin and 0.34 in fibroblasts. (d, e) Lesional skin; constrained and tolerant genes are the most and least loss-of-function-intolerant LoFtool deciles (1,237 and 1,119 genes with complete architecture data). (d) Observed over expected private coding variants: 1.12 in constrained and 1.06 in tolerant genes (ratio 1.05, P = 0.20). (e) Missense and truncating variants relative to synonymous variants within each class, against the expectation from site composition (95% CI); P refers to the constrained/tolerant ratio for missense variants. Constrained/tolerant ratio: missense 0.75 (0.63–0.89, P = 0.0008), truncating 0.76 (0.56–1.04, P = 0.083; 108 events). Depletions below 0.90 (missense) and 0.81 (truncating) were detectable.

We next tested for gene-level selection with dNdScv, which compares protein-altering with synonymous mutations while accounting for the sequence context of possible substitutions [27]. No gene showed significant positive selection in either skin or fibroblasts, and the global ratios for missense and nonsense mutations were close to one, consistent with near-neutral evolution. Per-gene binomial tests were likewise negative in skin and identified only one gene in fibroblasts. Simulation analysis showed that recurrent truncating mutations of modest frequency would usually have been detected, arguing that the negative result was not simply due to lack of power.

We next tested whether mutations preferentially accumulated in functional pathways. Screening 8,579 pathway terms against architecture-matched random gene sets identified two enriched categories in skin, extracellular matrix (ECM)-receptor interaction and the mitotic spindle hallmark, whereas none was significant in fibroblasts. Direct testing with the two calibrated exposure models weakened these signals: the ECM-receptor set remained enriched under one model and was only marginal under the other, while the mitotic-spindle signal was model-dependent. The ECM-receptor excess also lost significance after removal of individual influential genes, and synonymous variants were at least as enriched as missense variants, favouring locally elevated mutation rates over selection for altered protein function.

Because chromatin-modifying genes were prominent among the recurrently mutated genes in our previous study [2], we tested a prespecified panel of 69 such genes. In skin, this panel showed an excess of variants under the opportunity-spline model but not when callability was estimated freely, and no excess was present in fibroblasts. The apparent skin enrichment involved synonymous as well as protein-altering variants and was not driven by a single gene, again suggesting increased mutational opportunity rather than selection for altered chromatin-regulator function.

An alternative possibility is that protein-disrupting mutations in genes essential for cell survival or function are depleted by purifying selection [13,27]. We therefore compared genes with high and low intolerance to loss of function, ranked by LoFtool [28], using synonymous variants within each group as an internal control for gene length, sequencing depth, capture and local mutation rate. Overall mutation burden was similar between the groups (Fig. 5d), but missense variants were modestly depleted relative to synonymous variants in constrained genes, while truncating variants showed a similar but non-significant trend (Fig. 5e). Purifying selection was therefore detectable in SSc skin but removed only a minority of potentially damaging variants, and clones carrying damaging variants were not smaller than other clones, which is consistent with findings in cancer and in normal somatic tissues [13,27]. Damaging variants in DNA-repair, chromatin or oxidative-stress genes were not associated with a higher C>A fraction, so the mutational process could not be explained by somatic loss of DNA-repair function. In summary, the clones detected in SSc skin showed neither gene-nor pathway-level positive selection, and only weak purifying selection against damaging variants.

### 3.6. Somatic variants accumulate with age as small clones

To determine whether somatic variants accumulate over time in SSc dermis, we related private coding variant burden to age across patients aged 24 to 71 years. Disease duration was not associated with variant burden, whereas mutations increased steadily with age, also after adjustment for disease duration (Fig. 6a, b). All major variant classes contributed proportionally, consistent with a general age-related mutational process rather than selection for a particular protein consequence.

**Figure 6.**
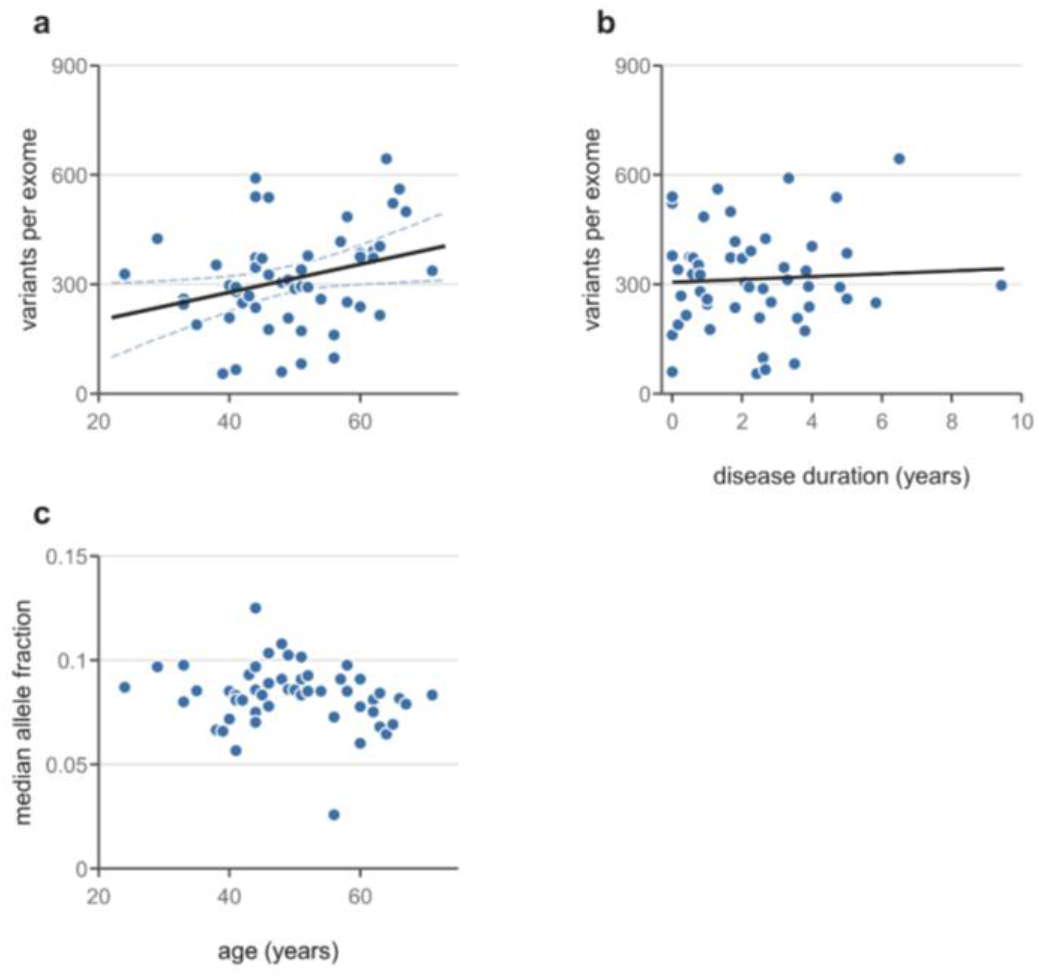
Somatic variants accumulate with age but not with disease duration. Fifty-two lesional skin exomes from 48 patients aged 24 to 71 years; private coding variants with at least three alternate reads unless stated. (a) Burden against age with least-squares fit and bootstrap 95% band (dashed); +3.85 variants per exome per year (95% CI 0.39–7.32), Spearman ρ = 0.31, P = 0.024, and P = 0.036 after adjustment for disease duration. The correlation was similar or stronger as weakly supported calls were excluded. (b) Burden against disease duration (ρ = −0.01, P = 0.96; median duration 2.1 years); the smallest effect detectable with 80% power was 27.7 variants per exome per year. (c) Median allele fraction per exome against age (ρ = −0.17, P = 0.24). Small clones (allele fraction below 0.05) increased by 2.66% per year (95% CI 1.11–4.21) and larger clones (0.05–0.35) by 0.43% per year (−0.24 to 1.08) in negative binomial models adjusted for median depth and callable coding sequence with patient-clustered errors.

We next asked whether the greater mutation burden in older patients reflected expansion of pre-existing clones or accumulation of new, small clones. Median allele fraction was unchanged with age (Fig. 6c), whereas low-frequency variants became more common. After adjustment for sequencing depth and callable coding sequence, the number of small clones increased with age by approximately 3% per year, whereas larger clones showed no age-related expansion (0.4% per year, not significant). The age-associated increase in mutation burden therefore reflected the appearance of additional small clones rather than progressive growth of established ones.

Taking into account the relationship between mutation burden and patient age (Fig. 6a), we considered the possibility that the somatic landscape reflected fibroblast senescence rather than the ongoing genotoxic stress suggested by our analyses in Figure 3. Within the expression programme carrying the length-dependent transcriptional loss, non-secreted SenMayo senescence-associated genes showed no association, while secreted senescence-associated (SASP) genes tracked the difference between SSc and normal fibroblasts but not variation within either group (Fig. 3f). Growth-arrest genes followed the programme, but proliferation genes were not suppressed, indicating that the affected fibroblasts were not in a canonical growth-arrested state. SSc-associated expression changes were also essentially unrelated to known age-associated transcriptomic changes [29], and genes annotated as pro- or anti-longevity [30] were mutated at the rate expected from their mutational opportunity. In summary, these findings favour a cumulative effect of persistent cellular stress over premature ageing or classical senescence as the setting for somatic mutagenesis in SSc fibroblasts.

### 3.7. Somatic clones persist after autologous stem cell transplantation

Because AHSCT improves fibrosis and overall survival in dcSSc [31–33], we tested whether clinical remission depended on elimination of mutation-bearing dermal clones, using 10 lesional-skin and 12 fibroblast exomes from 11 patients obtained 2 to 33 months after AHSCT, including seven paired pre- and post-transplant samples from four patients.

After adjustment for age, skin from AHSCT recipients and from untreated patients showed no detectable difference in coding variant density, C>A fraction, dominant-clone density or median allele fraction, and none of these measures varied with time since transplantation (Fig. 2f). Because some conditioning regimens include total-body irradiation, which can generate small deletions [34], we also examined indels separately and found no post-transplant increase; total-body irradiation had been given to only a minority of the patients in this group.

In the seven paired samples, four variants were detected both before and after transplantation. No shared variant was found in any of 36 mismatched pre- and post-transplant patient pairs, indicating that the repeated variants marked patient-specific clones that persisted through a conditioning regimen containing high-dose cyclophosphamide [32,35] rather than recurrent artefact. All four were non-coding and weakly supported, so they establish persistence of clones only tentatively. In summary, clinical remission after AHSCT was not accompanied by elimination of mutation-bearing dermal clones.

### 3.8. Somatic mutations in autoantigen genes rarely create neo-epitopes

Somatic mutation can generate neoantigens capable of initiating immune responses, as demonstrated by immunity against RNA polymerase III in SSc patients with coincident cancer harbouring mutations of *POLR3A* [36]. We therefore asked whether somatically mutated dermal cells could provide an analogous source of altered autoantigens by examining coding variants in prespecified panels of connective-tissue-disease autoantigens.

Coding mutations in autoantigen genes were found in a substantial minority of samples. In fibroblast cultures, the most frequently affected genes included several RNA polymerase subunits, *POLR1A*, *POLR2A*, *POLR1B* and *POLR3A*, as well as *PRKDC*, and several of these genes were also mutated in skin (Fig. 7a). However, when compared with 5,000 random gene sets matched for mutational opportunity, both the core 18-gene panel and the extended autoantigen panel were mutated at approximately the expected background rate in skin and fibroblasts, and none of the 28 antigen-specificity groups showed significant enrichment. Autoantigen genes were therefore not preferentially targeted by somatic mutation.

**Figure 7.**
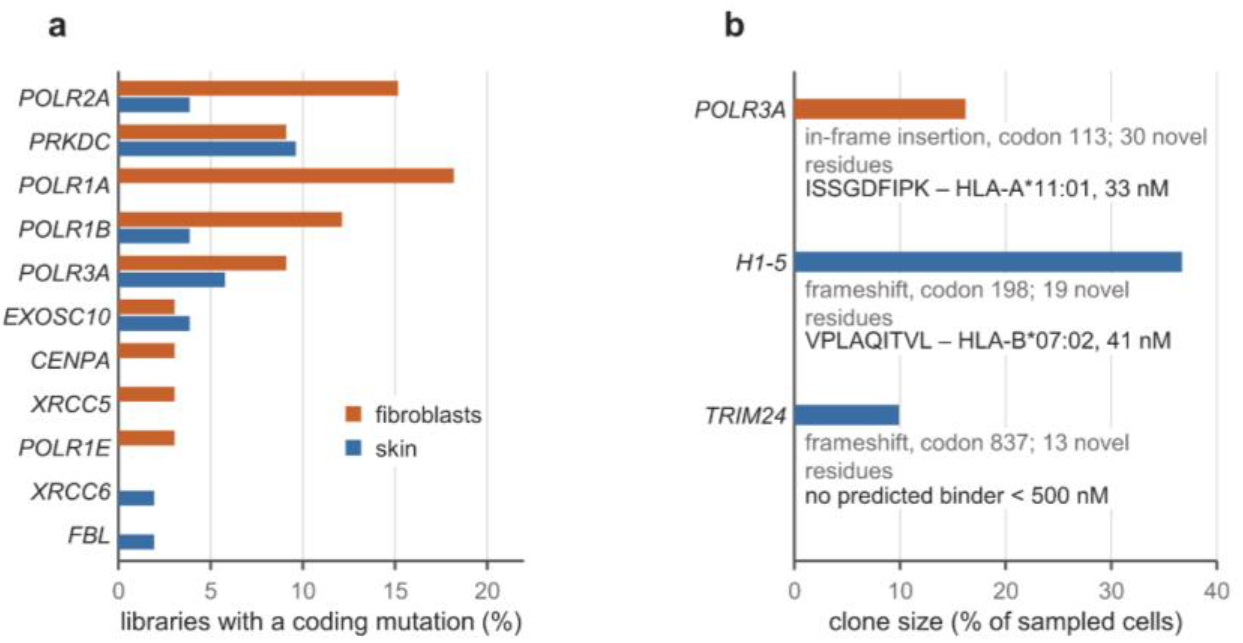
Somatic muta-tions in autoantigen genes. (a) Percentage of libraries with at least one coding mutation in genes of the prespecified 18-gene SSc autoantigen panel, in 52 skin and 33 fibroblast exomes; the seven panel genes without any coding mutation are not shown. (b) The three mutations that introduce amino acids absent from the germline proteome, with the clone size of the carrying variant (*POLR3A* 16%, *H1-5* 37%, *TRIM24* 10% of sampled cells), the number of novel residues and the strongest predicted class I binder from each novel stretch (MHCflurry 2.2.1 [37], 14 population-representative alleles).

We next asked whether protein-altering mutations in autoantigen genes generated genuinely novel sequences. Only three of 135 variants did so: a 30-residue in-frame insertion in *POLR3A* in a fibroblast culture and frameshifts in *H1-5* and *TRIM24* in skin, producing novel C-terminal sequences carried by sizeable cell clones (Fig. 7b). All other frameshifts terminated within one or two codons. Across 14 population-representative HLA class I alleles, the *POLR3A* insertion and the *H1-5* frameshift each generated a peptide predicted to bind strongly without a wild-type counterpart, whereas the *TRIM24* frameshift did not. Likewise, only a small minority of missense peptides gained predicted HLA binding relative to their wild-type sequence. None of the three carriers had a recorded malignancy, and anti-RNA polymerase III and anti-TIF1α serology were negative, which prevented direct assessment of an immune response to the altered sequence. Thus, with the possible exception of the *POLR3A* insertion, somatic mutations in dermal cells were unlikely to provide a primary source of neo-epitopes in SSc.

## 4. Discussion

The principal finding of this study is that lesional dermis from patients with SSc contains a substantial burden of somatic mutations, including a C>A component associated with the severity and duration of skin disease. Weighted by the proportion of cells carrying each variant, untreated SSc dermis contained 5 to 8 coding mutations per megabase per cell, several-fold more than reported in healthy skin fibroblasts and other normal tissues [38–42].

Several observations support oxidative damage as the source of these C>A mutations. The substitutions lacked transcriptional strand asymmetry, were depleted at CpG sites and were enriched in thymine-preceded contexts, which is consistent with 8-oxoguanine mispairing rather than with helix-distorting DNA adducts. Our interpretation agrees with previous evidence of constitutive reactive oxygen species production in SSc fibroblasts [4], of DNA damage and chromosomal instability [8], and of increased γH2AX foci in diffuse disease [6]. Systemic oxidative injury has also been demonstrated through DNA damage in peripheral blood mononuclear cells [43] and increased urinary 8-oxo-2′-deoxyguanosine associated with pulmonary involvement and skin fibrosis [5]. Our findings extend these observations from markers of damage to its mutational consequence, showing that the C>A fraction, read as an index of cumulative oxidative injury, correlated with damage accrual in the skin as estimated by the product of modified Rodnan skin score and disease duration. This is distinct from SSc-associated interstitial lung disease, where mutagenesis was attributed primarily to translesion polymerase and deaminase-like activity [3].

Transcriptional data from fibroblasts provided further evidence of persistent genotoxic stress. DNA lesions stall elongating RNA polymerase II and reduce transcription in proportion to the length of DNA traversed, so that genes with long transcription units are preferentially affected [19]. Consistent with this mechanism, SSc fibroblasts showed progressively reduced expression with increasing intronic length, and the programme carrying that decline was associated with ATF4-dependent integrated stress-response genes, consistent with our demonstration of PERK-ATF4-FOXO1 activation in SSc fibroblasts [6]. DNA damage may also be linked to increased *ATF4* activity and to the translation of particular mRNA species through N⁶-methyladenosine modification, which regulates both the DNA damage response and *ATF4* transcriptional initiation [44–46].

Somatic mutations occurred in expanded cellular clones, with detected variants typically present in approximately one fifth of sampled cells. However, neither gene-level nor pathway analyses identified convincing positive selection, and purifying selection against damaging variants was weak. This contrasts with normal epithelia, where clones carrying cancer-associated driver mutations frequently undergo positive selection [12,13,47]. Clonal expansion in SSc therefore appears largely independent of the functional consequences of individual mutations. Expansion nevertheless implies that mutation-bearing cells can survive and divide despite genotoxic stress, and we have recently shown that dcSSc fibroblasts resist mitochondrial apoptosis, which can be restored by inhibiting PERK or FOXO1 [6]. Although senescence-associated markers are increased in SSc [6,10,48], our recent transcriptomic analysis identified predominantly a secretory senescence-like programme without suppression of proliferation-associated transcriptional changes [21]. These transcriptional observations stand in contrast to the reduced cell numbers and reduced Ki67 labelling of dcSSc fibroblasts in culture (Khan et al. 2026), and the discrepancy is not resolved by the present data. One possibility is that the mutation clones we detected divided earlier in the disease and have since slowed, since clonal expansion records past division rather than current proliferative rate. Enrichment of mutations in late-replicating DNA [24] likewise places the mutational process in dividing rather than arrested cells. Senescence-associated signals in dcSSc fibroblasts may therefore reflect an imbalance between accumulating replication-associated DNA damage and protective responses driven by STING and by PERK-ATF4 dysregulation, with somatic mutation as one durable consequence.

AHSCT provides an opportunity to examine whether mutant clones are required to maintain fibrosis. We have recently shown that, compared with dcSSc fibroblasts, fibroblasts obtained after AHSCT carry reduced signals of genotoxic stress, reduced senescence-associated signals and restored sensitivity to mitochondrial apoptosis [6]. In the present study, mutation burden, dominant-clone density, allele fractions and the C>A component of post-AHSCT samples remained comparable to those of untreated dcSSc. Regression of fibrosis therefore did not require elimination of detectable mutation-bearing clones, which argues that their persistence alone is insufficient to maintain the fibrotic phenotype. Somatic mutations may instead persist as a record of previous injury, although a contribution of surviving clones to later relapse cannot be excluded, and relapse or progression occurs in approximately one third of transplanted patients [35,49].

Somatic mutations have also been proposed as a source of autoantigens in SSc. In patients with anti-RNA polymerase III antibodies, temporally associated malignancies can harbour *POLR3A* mutations that elicit mutation-specific immunity and cross-reactive antibodies [36,50]. We found no enrichment of mutations in autoantigen genes beyond their expected mutational opportunity, and only three of 135 protein-altering variants generated novel amino acid sequence, including a POLR3A insertion supported by three sequencing reads in a fibroblast culture from a patient without anti-RNA polymerase III antibodies. These findings provide no compelling evidence that somatic mutations in dermal cells initiate SSc autoimmunity, although mutations in unsampled tissues cannot be excluded, particularly given the restricted autoantibody repertoire and subsequent epitope spreading characteristic of SSc [51].

Our findings differ from those in SSc lung fibroblast lineages, where kataegis, polymerase η- and deaminase-like signatures, and structural genomic alterations were reported [3]. Tissue-specific biophysical properties, such as the higher oxygen tension in the lungs compared with the skin, may contribute to these differences. This is relevant to senescence-associated signals, because mild hyperoxia accelerates telomere shortening and blocks fibroblast proliferation in vitro [52]. Telomere shortening has been associated with progression of SSc-associated lung disease but not with skin disease [53], while anti-TERF1 antibodies and germline telomerase defects are linked to pulmonary fibrosis [54,55]. Direct comparisons across affected organs will be needed to establish whether distinct mutational processes operate in different SSc tissues.

Several limitations should be noted. Detection of low-frequency variants by bulk exome sequencing, which remains susceptible to library-preparation and sequencing errors, particularly oxidative C>A artefacts [56]. The consistency of the mutational patterns across samples and clinical groups argues against artefact as the sole explanation, but rare variants and small clones require confirmation using higher-fidelity approaches such as duplex sequencing or whole-genome sequencing of single-cell-derived clones [13,41,42]. Chromatin accessibility was not profiled, so the relationship between open and closed chromatin and the distribution of somatic variants was approached only indirectly through replication timing and transcriptional output.

In summary, the somatic genome of SSc skin appears to reflect two overlapping processes: predominantly neutral mutations accumulating with age, and a C>A-rich component associated with cumulative skin disease and compatible with persistent oxidative DNA damage. Mutation-bearing cells can survive and expand without detectable positive selection, leaving a durable genomic record of tissue injury that persists even after clinical regression of fibrosis.

## 5. Methods

### Study approval

The study was approved by the University of Alberta Research Ethics Board (Pro00085583, Pro00090050). All participants gave written informed consent before sampling, in accordance with the Declaration of Helsinki.

### Patients and samples

Lesional skin biopsies were obtained from patients attending the systemic sclerosis clinic at the University of Alberta, embedded in OCT, and microdissected to fibrotic dermis. Fibroblast cultures were established from the same biopsy and grown to passage 2 in Dulbecco’s Modified Eagle Medium (DMEM, Thermo Fisher, cat# 211965118) with high glucose containing L-glutamine, 1% penicillin-streptomycin, and 10% fetal bovine serum (FBS, Thermo Fisher cat# 12484028). Buccal swabs served as the germline reference. The analysed set comprised 85 exomes from 55 patients: 52 lesional skin exomes from 48 patients (42 untreated and 10 obtained after AHSCT) and 33 fibroblast exomes from 29 patients (21 untreated and 12 post-transplant); 22 patients contributed both tissues. Treatment status was assigned at the time of each biopsy. Clinical variables (age, sex, disease duration, time since transplantation, modified Rodnan skin score, organ involvement, smoking history coded as never against ever, autoantibody serology and the Scleroderma Clinical Trials Consortium Damage Index) were taken from the clinical record.

### Sequencing, variant calling and quality control

Exomes were captured with SureSelect Human All Exon and UTR V6, Agilent Technologies (91 Mb padded design) and sequenced on Novaseq X Plus, 25B read flow cell. Reads were trimmed with fastp v1.3.5, and then processed following the GATK Best Practices data pre-processing workflow: alignment to GRCh38 (Broad hg38 reference, Homo_sapiens_assembly38) with BWA-MEM v0.7.19, duplicate marking and base quality score recalibration (GATK v4.6.2.0). Somatic SNVs and indels were then called following the GATK Somatic short variant discovery workflow with Mutect2 (GATK v4.6.2.0), using matched buccal DNA as the normal control, together with a panel of normals and the gnomAD germline resource. Calls were filtered with FilterMutectCalls, using contamination estimates (GetPileupSummaries / CalculateContamination) and read-orientation artifact priors (LearnReadOrientationModel). Consequences were annotated with VEP including LoFtool and Condel scores [28,57]. Nineteen libraries were excluded before analysis: ten skin libraries without a usable matched-normal call set (four with failed buccal sequencing) and, by design, one healthy-control fibroblast line and the tissue-exome series. Four further libraries were excluded because more than 30% of their private calls lay between allele fractions of 0.35 and 0.65, the window of germline heterozygous sites (flagged libraries 0.31–0.81, next-highest 0.24); every excluded library had a passing library from the same patient. Only sites observed in a single patient (private calls) were analysed. Every analysis was repeated at minimum alternate-read thresholds of one, three, five and eight; three reads is the primary threshold. Because the fibroblast C>A fraction was elevated only among weakly supported calls and reached the skin value at eight reads, analyses of the C>A component were restricted to skin.

### Callable territory and expected counts

Per-gene callable bases at 10-fold depth or more were computed per library with mosdepth [58]. Mutational opportunity per gene was taken from RefCDS (GRCh38) [27], multiplied by the captured fraction of coding sequence and by the exact callable fraction (mean callable bases across libraries divided by captured territory). Difficult-region and low-mappability fractions were taken from the Genome in a Bottle (Genome in a Bottle | NIST) stratifications. Gene-set excess was estimated with negative binomial models (dispersion θ = 3.6) in two forms calibrated on 2,000 architecture-matched null sets: an offset in callable opportunity with a natural spline in log opportunity, and a capture-only offset with free splines in coding length and callable fraction.

### Spectrum, context and signatures

Substitutions were reduced to six pyrimidine-referenced classes and 96 trinucleotide channels from the coding-strand context of each variant, and per-library spectra were averaged within and then across patients. Context and strand enrichments were computed against per-gene, per-trinucleotide opportunity derived from RefCDS coding sequences, so the expected coding-strand fraction is the Poisson-binomial mean rather than 0.5. Signatures were compared by cosine similarity with COSMIC v3.4 single-base-substitution signatures rescaled to exome trinucleotide composition [59].

### Clone size and detection

Allele fractions were recomputed from alternate and total read counts. Clone size is the fraction of sampled cells carrying a variant, taken as twice the allele fraction under heterozygosity; clone-size analyses used private coding calls at 30-fold depth or more and allele fractions below 0.35. Small clones occupy less than 10% of sampled cells (allele fraction below 0.05) and dominant clones at least 40% (0.20 or above). Detection power is the binomial probability of at least three alternate reads at a given depth and allele fraction. The latent allele-fraction distribution was estimated by maximum likelihood with each call modelled as a binomial draw at its own depth, conditioned on detection, for log-normal, gamma, truncated power-law and beta families.

### Selection

Global and gene-level dN/dS were estimated with dNdScv [27] without covariates, separately for skin and fibroblasts, on private calls with alleles of 50 bp or shorter. Per-gene binomial tests against opportunity and against synonymous counts were prespecified at each read-support threshold, and power was estimated by adding simulated truncating mutations to 30 random genes. Mutation-dense genes (observed over expected at least 1.5 and at least five variants) were tested for over-representation in 8,579 GO, Reactome, KEGG and MSigDB hallmark terms against 1,000 stratum-matched permutations with Westfall–Young familywise control. Prespecified chromatin-modifier, spliceosome, DNA-repair and autoantigen panels were tested against opportunity-matched random gene sets with leave-one-gene-out refits. Constrained and tolerant genes were the lowest and highest LoFtool deciles, and damaging variants were nonsense, essential splice-site and Condel-deleterious missense variants.

### Transcriptomic and replication-timing analyses

Libraries from total RNA from 26 fibroblast cultures matching 27 exomes were prepared with Kapa RNA HyperPrep kit (Roche) and sequenced Novaseq X Plus, 25B read flow cell, Illumina. Reads were trimmed with fastp v1.3.5 and quantified with RSEM v1.3.3 [60], using Bowtie2 v2.5.5 as the aligner [61], against the GRCh38 reference with Ensembl release 113 annotation. Genes at one transcript per million or more were counted as expressed. Replication timing was taken from the six-fraction ENCODE Repli-seq profile of BJ fibroblasts (ENCFF001GIC, -GIP, -GIX, -GJC, -GJH, -GII) [25], averaged over the 100 kb window containing each gene with weights from 0.917 (G1b) to 0 (G2). Variant density was modelled per library by negative binomial regression with a log callable-length offset and adjustment for GC content, CpG density and difficult-region and low-mappability fractions, and coefficients were combined across patients. Fibroblasts were aggregated to pseudobulk log counts per million by donor and subtype; the SSc-minus-normal fold change was decomposed exactly over the principal components of the unit-by-gene matrix, and length-matched module scores subtracted the mean z-score of 50 control genes per member matched on genomic-span decile and expression quintile. External ageing references were the per-gene age correlations in mouse skin [29], mapped through MGI one-to-one orthologues, and GenAge model-organism longevity genes mapped to human orthologues [30,62].

### Clinical associations, age and transplantation

Age effects on burden and on small- and larger-clone counts were estimated with negative binomial models adjusted for median depth and callable coding sequence. Subtype and severity analyses used the 41 untreated early-disease skin exomes and binomial models of C>A against non-C>A counts; mRSS and the accumulated index (mRSS × disease duration) were standardised. A grid of four somatic measures against 13 clinical variables (48 testable combinations, one observation per patient) was fixed before analysis and corrected by Benjamini–Hochberg, with a block-permutation null of 2,000 replicates. Post-transplant exomes were compared with untreated early-disease skin in age-adjusted models. Persistence was assessed as private variants of any read support shared between pre- and post-transplant samples of a patient, against all 36 mismatched combinations of one patient’s first with another patient’s second sample.

### Autoantigen analyses

Coding mutations were examined in a prespecified 18-gene SSc autoantigen panel and an extended connective-tissue-disease panel grouped into 28 antigen specificities, against 5,000 random gene sets matched on mutational opportunity. Novel protein sequence was defined as amino acids absent from the germline proteome, and 8- to 11-mer peptides from novel stretches and from missense mutations with their wild-type counterparts were scored with MHCflurry 2.2.1 [37] across 14 population-representative HLA class I alleles; strong binders had a presentation percentile of 0.5 or below.

### Statistics

Multiple libraries from one patient are not independent, so per-library estimates were averaged within and then across patients, or standard errors were clustered by patient, or intervals were obtained by patient-level bootstrap. Tests were two-sided. Multiplicity was controlled by Benjamini–Hochberg within analysis families and by Westfall–Young familywise error for the pathway screen. Negative results are reported with the effect detectable at 80% power. Analyses used Python 3.11 (numpy, pandas, scipy, statsmodels) and R (dndscv).

## Author contributions

Conceptualization: R.G., M.O. Methodology: R.G., D.H., AI, S.O’K., M.O. Investigation: R.G., A.I., S.O’K., J.W. Resources: J.S., J.W., C.D., M.O. Formal analysis: R.G., D.H. Writing – original draft: R.G., M.O. Writing – review and editing: all authors. Supervision: R.G., M.O.

